# Effects of Neurofeedback training on performance in laboratory tasks: A systematic review

**DOI:** 10.1101/2022.10.14.511990

**Authors:** Payton Chiasson, Maeve R. Boylan, Mina Elhamiasl, Joseph M. Pruitt, Saurabh Ranjan, Kierstin Riels, Ashish K. Sahoo, Arash Mirifar, Andreas Keil

## Abstract

Neurofeedback procedures are attracting increasing attention in the neuroscience community. Based on the principle that participants, through suitable feedback, may learn to affect specific aspects of their brain activity, neurofeedback has been applied to basic research, translational, and clinical science alike. A large segment of the extant empirical research as well as review articles have focused on the extent to which neurofeedback interventions affect mental health outcomes, cognitive capacity, aging, and other complex behaviors. Another segment has aimed to characterize the extent to which neurofeedback affects the targeted neural processes. At this time, there is no current systematic review of the effects of neurofeedback on healthy participants’ performance in experimental tasks. Such a review is relevant in this rapidly evolving field because changes in experimental task performance are traditionally considered a hallmark of changing neurocognitive processes, often established in neurotypical individuals. This systematic review addresses this gap in the literature using the PRISMA method, building on earlier reviews on the same topic. Empirical studies using EEG or fMRI to alter brain processes linked to established, well-defined cognitive and affective laboratory tasks were reviewed. Substantial variability was found regarding the nature of the control for placebo effects, the implementation of the feedback, and the neural targets of feedback. Importantly, only a minority of the studies reported statistically meaningful effects of neurofeedback on performance in cognitive and affective tasks. Examining effect sizes and p-values in a subset of studies found no evidence for reporting bias, while also not finding systematic relations between study characteristics such as sample size or experimental control on the one hand and efficacy on the other. Implications for future work are discussed.

## Introduction

Neuroimaging techniques such as electroencephalography (EEG) and functional magnetic resonance imaging (fMRI) have provided key contributions to understanding neural processes in the human brain associated with specific aspects of behavior and experience. Increasingly, imaging techniques are also used in the context of intervention, often taking the form of neurofeedback training (NFT; Rogala et al., 2016; Thibault et al., 2018). This technique, also referred to as closed-loop training, relies on training participants to modulate their brain activity. To this end, NFT involves three steps: (i) measuring the targeted brain activity, (ii) presenting an index of the targeted brain activity to the participant (feedback), and (iii) enabling the participant to attain operant control over the measured activity through a suitable training regimen (Viviani & Vallesi, 2021). Despite these shared principles, studies using NFT vary strongly concerning how the three steps are implemented. Strong variability also exists regarding how the effects of NFT interventions are controlled through appropriate study designs, and regarding the goals of the NFT (Mirifar et al., 2022). Furthermore, pronounced inter-individual variability exists between participants, with significant proportions of non-responders frequently reported (Alkoby et al., 2018; Weber et al., 2020).

Paralleling neuroimaging techniques more broadly, NFT in particular, is used in basic research as well as in clinical and translational research (Sitaram et al., 2017). The majority of published research with NFT examines the effects of modulating one’s brain activity on clinical outcomes such as mental health challenges or neurological diagnoses (Arns et al., 2017; Begemann et al., 2016). However, basic science studies with healthy adults, targeting behavior as measured in well-established standardized tasks, are widely considered a gold standard for establishing the validity of psychological or neural interventions (Morris et al., 2022). Accordingly, establishing the specific mechanism of action and the nature of the evidence supporting an intervention technique will typically precede its clinical application.

Since its inception, questions have been raised regarding NFT’s efficacy and effectiveness (Schabus, 2017). A hallmark of an intervention being effective is its causal impact on its treatment target. In the case of basic science NFT studies, the goal of the intervention is not to improve symptoms in clinical groups, but to improve behavioral performance in non-clinical, healthy, samples (Gruzelier, 2014a). Often, these studies aim to test mechanistic hypotheses by means of using NFT to establish causal effects of certain brain processes concerning specific behaviors. For example, if down-regulating alpha power heightens the attentive selection accuracy for targets among distractors, this supports the hypothesis that EEG alpha power is part of a causal nexus linked to selective attention. Consequently, an obvious criterion of the effectiveness of NFT procedures is the extent to which NFT alters the target behavior, operationalized as performance in a defined laboratory task. A sizable body of previous empirical work and several review articles have examined this question (Mirifar et al., 2022; Sitaram et al., 2017). Collectively, this work however has not identified robust evidence for NFT’s effectiveness, sometimes concluding that NFT does not affect performance in psychological tasks (e.g., Rogala et al., 2016). Other authors however have reported robust outcomes, based on selective reviews of the literature (e.g., Gruzelier, 2014).

Previous review articles have also discussed several conceptual and methodological issues that vary widely in the published literature, including the number of training sessions, the usage of suitable control conditions, and the neuro-mechanistic framework used for deriving hypotheses (Gruzelier, 2014b; Rogala et al., 2016). More recent work has attempted to address these challenges and has increasingly used more sophisticated methods both in the implementation of NFT as well as in the experimental control of NFT (Watanabe et al., 2017). For example, multivariate analysis methods, often based on computational tools such as pattern classifiers or decoders, have become more widely used (Taschereau-Dumouchel et al., 2021).

Several questions however remain: (i) Across studies in non-clinical samples, is there evidence today for robust effects of NFT on performance indices of laboratory tasks? (ii) Is there a difference in outcomes between fMRI-based compared to EEG-based NFT methods? (iii) Is there a relation between finding evidence for altered performance and factors such as the brain imaging methods used (here: EEG vs. fMRI), the sample size, number of training sessions, or the targeted brain process?

The present review of the NFT literature published in the past 10 years examines these questions using the Preferred Reporting Items for Systematic Reviews and Meta-Analyses (PRISMA) approach. This approach ensures a systematic inclusion of published works to remove potential selection biases.

## Methods

### Study inclusion and PRISMA protocol

As shown in the PRISMA flow chart (Figure 1), a primary search was conducted on PubMed, complemented by additional searches on Google Scholar and PsychInfo. The following PubMed search terms and their equivalent on the other databases were used; For EEG-based NFT studies:

**Figure 1.**
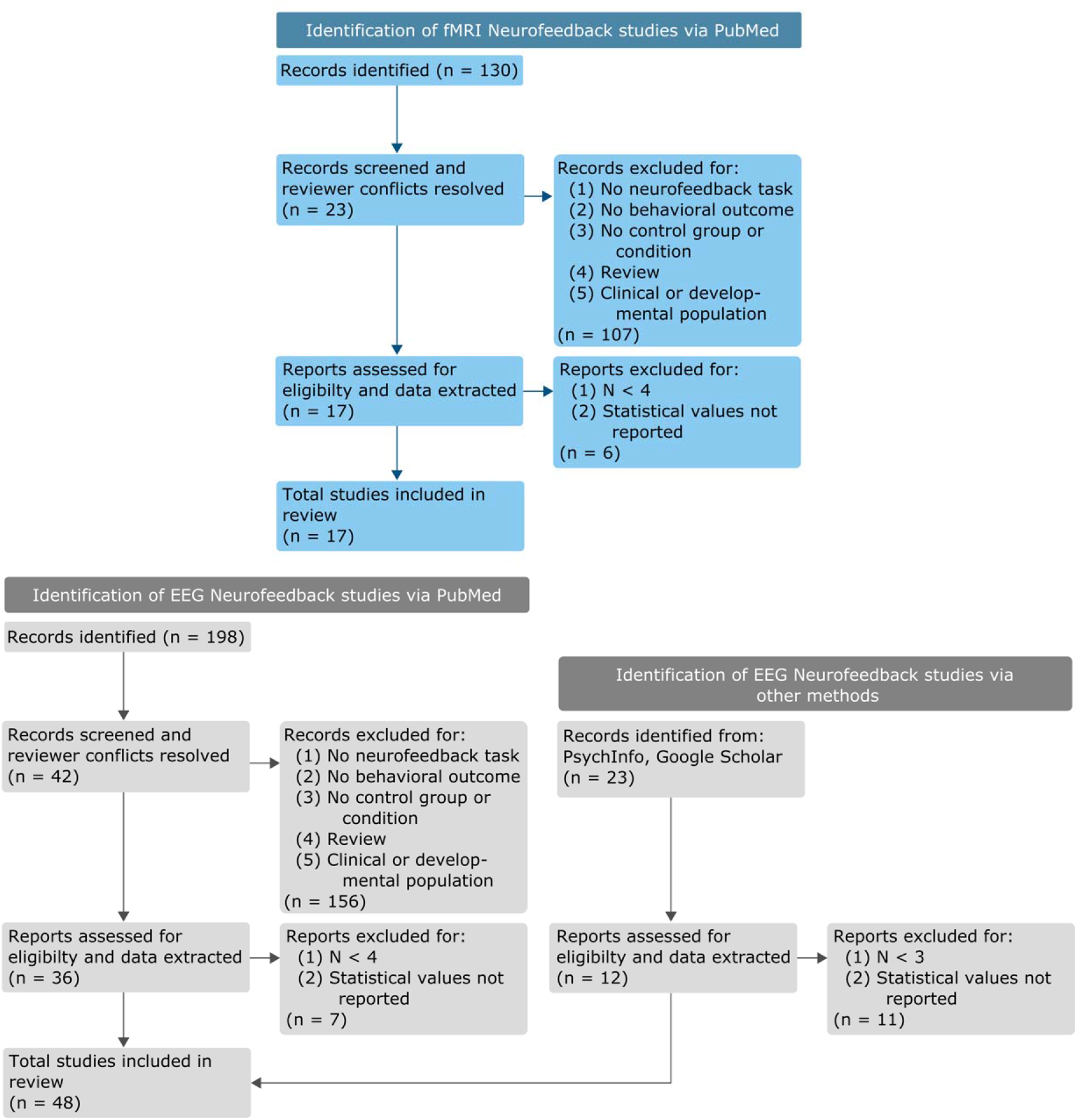
PRISMA review process of neurofeedback papers included/excluded for analyses. Top panel: PRISMA process for fMRI studies; Bottom panel: PRISMA process for EEG studies.

(((neurofeedback) AND (EEG)) AND (Task) AND ((“2012/01/01”[Date - Publication] : “3000”[Date - Publication]))) NOT Anxiety[Title] NOT Depression[Title] NOT Therapy[Title] NOT Disorders[Title] NOT psychiatric[Title] NOT treatment[Title] NOT review[Title]).

For fMRI-based NFT studies:

(((neurofeedback) AND (fMRI)) AND (Task) AND ((“2012/01/01”[Date - Publication] : “3000”[Date - Publication]))) NOT Anxiety[Title] NOT Depression[Title] NOT Therapy[Title] NOT Disorders[Title] NOT psychiatric[Title] NOT treatment[Title] NOT review[Title]).

These search terms aimed at identifying basic science studies (instead of studies targeting clinical diagnoses), which included psychological tasks, while avoiding review papers. The present review focused on published papers from the past 10 years (2012 to 2022) to capture studies with improved methodological standards such as the usage of appropriate control groups as recommended in seminal review papers published prior to and during the present inclusion period (e.g., Gruzelier, 2014b; Vernon, 2005). The initial search resulted in a total of 351 papers, including papers that were obtained from screening the reference sections of the search results.

These records were then screened by at least two reviewers, and excluded if they met the following exclusion criteria: (1) Exclude **Reviews** that do not present any new data; (2) Exclude Papers that have **neuro-atypical population;** (3) Exclude papers about **developmental** populations (children and older adults); (4) Exclude papers **without a behavioral** measure/outcome/metric; (5) Exclude papers with **no control** condition/ group; (6) Exclude papers published **before 2012**; (7) Exclude papers that **do not perform NFT**. Papers using a version of Brain-Computer Interfacing (BCI) with a robotic device were included if they involved a training phase in which feedback was given and a behavioral outcome was measured while meeting all the inclusion criteria above. Reviewer conflicts were resolved by adding at least one additional reviewer, followed by discussion, and vote.

Upon assessment and resolution of reviewer conflicts, this step resulted in a total pool of 87 published reports, 22 EEG fMRI, and 65 with EEG. These publications were then submitted to a systematic extraction process. In the process, it was found that a subset of the studies did not report inferential or descriptive statistics for the behavioral outcome. These studies were removed. Also removed were studies that reported inferential statistics such as ANOVA or t-tests with cells of N < 4. After this step, there were 48 studies with EEG and 16 studies with fMRI, included in the present report.

### Extraction of information from included papers

We extracted the information as shown in Tables 1 and 2 from each manuscript that entered the final sample. Specifically included were the type of experimental task used to measure behavioral outcomes, sample size, targeted brain response, age of the sample if indicated, type of feedback display, group design used for control, effect size, and the extent to which the main behavioral outcome variable(s) displayed a significant effect of NFT. This latter point proved challenging because many included studies provided incomplete or selective reports of statistical descriptors of behavioral outcome measures, as well as including several behavioral outcomes, quantified with several different indices each (e.g., response time, accuracy, d’, etc.). These cases were scored using the overall outcome utilizing the main variable mentioned in the introduction or by counting the number of significant versus null effects on relevant tests. Additional pieces of information that were often not reported in the original manuscript were participants’ handedness and duration of the experimental session. The nature of the control group or control condition were also extracted and categorized, combining active conditions (e.g., sham stimulation, yoked designs, feedback from control regions, or feedback from control EEG frequencies) and contrasting them with passive conditions (behavioral training only, waiting list).

**Table 1.** fMRI Neurofeedback Studies Included in the Present Review.

| Reference | Task | Sample size | fMRI target | Age | Feedback display | Design/control | Significant Effect |
| --- | --- | --- | --- | --- | --- | --- | --- |
| Habes et al., 2016 | binocular rivalry: two tasks categorization task and rivalry task | 17 | parahippocampal place area (PPA) over the fusiform face area (FFA) |  | thermometer* | no-NFT group in mock scanner | no |
| deBettencourt et al., 2019 | Memory recall | 24 (n<sub>eff</sub> = 18) | Based on two localizer runs, viewing blocks of scene, face, and object images. | M = 20.9 |  | Exp1: Feedback based on valid vs invalid/ Exp2: NF group vs yoked controls | yes |
| Oblak et al., 2021 | postneurofeedback motor task | 14 (n<sub>eff</sub> = 10) | motor cortex, primary sensorimotor cortex (M1 + S1) | M = 25.5, SD = 5.2 | grey disk | Within, baseline versus two feedback methods (ring vs middle) | yes |
| Scheinost et al., 2020 | not-XCPT/SART; AXCPT; VSTM; MOT | 25 (n<sub>eff</sub> = 20) | connectome | M = 21.5 | gas gauge | yoked control*sham | no |
| Herwig et al., 2019 | stimulation with emotional pictures | 40 (24/15 nft, 16/11 control) | amygdala with the anterior cingulate cortex, hippocampus, and distinct prefrontal areas | control M = 27.3, SD = 7.3, nft M = 26.7, SD = 4.8 |  | the control group also trained emotion regulation but without obtaining feedback | yes |
| Cortese et al., 2017 | motion discrimination task | 17 (n<sub>eff</sub>=10) | inferior parietal lobule (IPL), inferior frontal sulcus (IFS), middle frontal sulcus (MFS), and the middle frontal gyrus (MFG) | for 10 participants - M = 24.2, SD = 3.2 |  | for MVPA: motion discrimination task with 2-alternative forced choice discrimination, for NF training down-up group (first induce low, then high confidence) and vice-versa (up-down) | yes |
| Al-Wasity et al., 2021 | Go/No-Go | 20 (t= 10/5f(26.1 ± 5.1 years); c = 10/3f(23.2 ± 2.6 years) ) | SMA (NF task: User's choice MI of own execution of complex actions) |  | thermometer | true Nf vs sham NF | yes |
| Scharnowski et al., 2015 | Reaction time test, word memory task | 7 | Supplementary motor area and parahippocampal cortex | Range - 23-26y | continuously updated yellow curve | no control, three volunteers were trained to up-regulate in regulation blocks 1,3, and 5, whereas four volunteers up-regulated in regulation blocks 2,4, and 6. Also the type of feedback signal was pseudo-randomized: Three volunteers were trained to control the SMA-PHC feedback signal, and four volunteers were trained to control the PHC- | no |
|  |  |  |  |  |  | SMA feedback signal. |  |
| Wang et al., 2020 | PANAS |  | feedback signal based of the left amygdala (recalling past positive or pleasant events) |  | thermometer | true NF vs sham NF | no |
| Marins et al., 2019 | a motor imagery task, with no overt movement |  | fractional anisotropy (FA) in the sensorimotor segment of corpus callosum and connectivity of the sensorimotor resting state network |  |  | randomized, double-blind and sham-controlled | no |
| Paret et al., 2016 | subjective (affective) ratings of images | 32 (16 exp, 16 control) | amygdala or control region | 24.56 (3.91) | thermometer (continuously) | sham group, feedback from non-amygdala region | no |
| Kim et al., 2019 | questionnaire data | 60 | frontoparietal network, default mode network, salience network | 25.1 (2.9) | thermometer | sham group, feedback from other person | no |
| Chiew et al., 2012 | RT | 18 (13 exp, 5 control) | R & L M1 | 27 (3) exp, 24 (3) control | vertical bar | yoke sham group | no |
| Hui et al., 2014 | finger tapping accuracy | 28 (15 exp, 13 control) | PMA (pre-motor) | 22(1.6) exp, 23(1.7) control | continuously displayed feedback, green bar | control group | no |
| Li et al., 2016 | PANAS | 23 (12 exp, 11 control) | whole brain | 23.8 (1.4) exp, 23.1 (1.3) control | thermometer | control group | no |
| deBettencourt et al., 2015 | identification/attention task | 80 (32 in fMRI NFT, 48 in behav control) | uni: FFA, PPA; multi: "perceptual and attentional networks" | 20.3 | none | yoked control groups | yes |
| Wang et al., 2020 |  | 18 | V1/V2 | 23.4 ± 2.4 years |  |  | yes |

**Table 2.** EEG Neurofeedback Studies Included in the Present Review.

| Reference | Task | Sample size | EEG target | Age | Feedback display | Design/control | Significant Effect |
| --- | --- | --- | --- | --- | --- | --- | --- |
| Li et al., 2021 | laparoscopy | 20 | posterior electrode cluster |  | sphere*visual | no-NFT group* | yes |
| Sidhu & Cooke., 2021 | walking, serial sevens | 25 | Cz |  | tone pitch*auditory | crossover, up, down, sham*crossover design | yes |
| Hsueh et al., 2016 | working and episodic memory tasks | 50 | C3, C4, and Cz | ( NF: M = 20.96, SD = 2.85) / (Ctrl: M = 21.6, SD = 2.40) | horizontal bar | sham | no |
| Chang et al., | auditory | 12 | frontopolar midline |  | disk around | yoked control | yes |
| 2021 | discrimination and recognition tasks |  |  |  | fixation point |  |  |
| Arvaneh et al., 2019 | random dot motion task | 28 | Fz, C3, Cz, C4, P3, Pz, P4, and Oz(all channels) | (NF: M = 29.75, SD = 4.91)<br>(Ctrl: M = 25.57, SD = 3.50) | showed the identified letter in a text box | no-NFT group | yes |
| Wei et al., 2017 | backward digit span task and word-pair test | 30 | C3 | M = 26, SD = 3 | horizontal bar | sham | yes |
| Power et al., 2020 | unimanual button pressing task (but didn't report handedness, only used left-hand for button pressing) | 28 | C3 and C4 | M = 22, SD = 7 | bar | yolked control | yes |
| Karran., 2019 | business task (stock) | 30 | all electrodes | 18–43, $\mu$ 24 | color of task display frame | no-task, continuous NFT, event-related NFT | no |
| Eschmann & Mecklinger., 2020 | delayed match t samp, stroop | 35 (17 active 18 controls) | Fz | 20-30, $\mu$ 23.99 | | active control group (random frequency NFT) | no |
| Berger & Davelaar., 2018 | stroop | 22 (11 in each group) | Fp2 | 35.2, SD = 8.8 | floating object | 2-D VR and 3-D VR, no other control | no |
| Nan et al., 2020 | Continuous tracking task | 23 (12, 11 in each group) | C3 | 25.48 ± 3.62 years | bird chirps | NFT versus sham control | no |
| Mishra et al., 2021 | feature-based attention | 48 (32, NFT, 16 sham) | fonto-posterior source space | 25.5±0.3 | scale with plus signs | NFT vs Sham | no |
| Kim et al., 2021 | speech in noise detection | 20 | Fz, FCz, FC1, FC2, and Cz | 23.2 years; SD = 1.33 | plus sign | Placebo | no |
| He et al., 2020 | cued pinch movement | 20 | C3 C4 | 18-21 | basketball | sham | yes |
| Grosselin et al., 2021 | relax VAS | 48 (25, 23) | RMS | mean 33.3 range 18-60 | tone volume | sham | no |
| Agnoli et al., 2018 | alternative use task | 80 (40 alpha, 40 Beta, in each 20 sham 20 NFT) | CP2, CP4, CP6 and P4 | 21.10 years, SD = 2.12;(19 to 27 years) | natural picture stream progress | sham | no |
| Ros et al., 2013 | mind-wandering question, RT, and auditory oddball RT | 34 | Pz | 32.6, SD: 10.7 | dynamic bar graph | sham | no |
| Ozga et al., 2019 | mental rotation | 33 | 19-channels (all used for NFT) | 32.06; SD □ = □ 7.6 7 | 440 Hz tone, an intensity of 73 dB SP | no NFT | no |
| Ros et al., 2013 | sequence response time | 10 | C4 | 35.7, SD: 12.7 | start/stop movement of games | no NFT crossover 7 days apart | yes |
| Salari et al., 2014 | object detection and spatial att | 24 | PO7/PO8 | 29, range 25–35 | numbers and ball game | sham control | yes |
| Enriquez-Geppert et al., 2014 | task switch; 3-back; stop signal; stroop | 31 | Fz | 25 + -3 | display color -> red | sham control | no |
| Enriquez-Geppert et al., 2014 | task switch; 3-back; stop signal; stroop | 40 | Fz | 24.8 + -3.3 | display color -> red | sham control | yes |
| Xiong et al., 2014 | WM task | 48 (12 per group) | Fz, FCz, Cz, C1 and C2 | not reported | totating sphere and thermometer | only behav training, no trialing, sham | yes |
| Boe et al., 2014 | sequential button press paradigm | 18 | motor cortex (MEG) | 24.7 ± 3.8 | 2 bar graphs (l,R) | no NFT (fixation) control | no |
| Zich et al., 2017 | thumb of finger pinch movement (RT) | 20 | motor cortex (C3 or C4) |  | basketball | no training, sham | yes |
| Gordon., 2020 | packmen and ball filling | 140 | Pz |  | thermometer | silent control, active control | no |
| Renton et al., 2021 | perceived average motion direction | 30 | occipitoparietal |  | motion coherence |  | no |
| Yun et al., 2020 | controlling' the level of spontaneous audio-visual alpha band cortical connectivity. | 16 (22-40y) | temporal and occipital |  | bar graph | illusion (perceiving two flash) vs nonillusion (perceiving one flash) trials | yes |
| Parsons & Faubert., 2021 | 3D-MOT | 40 (m=22.89) | Pz |  | sphere/color saturation | non-active control, sham neurofeedback | yes |
| Faller et al., 2019 | boundary avoidance task (virtual flight) | 20 |  |  | loudness of audio of a constant-rate heart-beat. | sham, silence |  |
| Gonçalves et al., 2018 | Attention Network Task (ANT), WM: hought Identification Task (TIT), Content of WM: Resting State Questionnaire (ReSQ), MW Intentionality: Mind Wandering Deliberate and Spontaneous Scales (MW-D/S) | 30 (18 to 32) | Cz | M = 20.7, SD = 3.7 | bar graph | SMR↑Theta↓ Vs Theta↑SMR↓ | no |
| Reiner et al., 2018 | finger sequence task (average speed) | 45 | Pz | 28.26, s.d. 4.81 | bar graph | theta NFT, beta NFT, noNF | yes |
| Naros et al., 2016 |  |  |  |  |  |  | no |
| Holger Gevensleben1 et al., 2014 | continuous performance task/target detection | 20 | Cz | Exp: 23.2 (2.91)<br>Placebo: 22.9 (2.98) | ball on a computer screen | sham group | no |
| Escolano et al., 2012 | mental rotation task, psych battery, stroop, trail making test | 19 (10 NF, 9 sham) | P3, Pz, P4, O1 and O2 | NF 25.8 ± 4.07;<br>Control 24.3±3.67 | red or blue square | yoked sham | no |
| Schneider et al., 2019 | validity effect (VE) | 14 | PO7 and PO8 | 23 ± 1.52 | colored circles with an inscribed cross | repeated randomly assigned sham/NF training | yes |
| Reiner et al., 2014 | finger tapping | 38 | Pz | 25-35 | car acceleration | silent control (watched movies) | yes |
| Kober et al., 2015 | they learned to voluntarily increase SMR | 20 | P3 | M = 24.40 yrs., SE = 1.85 yrs. | scatter plots | received sham feedback | yes |
| Eschmann et al., 2021 | finger tapping | 46 | central sites, Fz, | Mage = 22.44, range = 18-30 years | bar charts | nonresponders were control (poor control) | yes |
| Eschmann et al., 2020 | source memory for words | 36 | central sites, Fz, | 23 | rollercoaster animation, | active control (random frequency) | no |
| Maszczyk et al., 2020 | simple and complex reaction time | 12 | C3 | 22-25 | airplane flying with sound | sham | yes |
| Nan et al., 2013 | peripheral visual feature conjunction detection | 30 | Cz | 19-33 | sphere and cube size and color | passive (no training) control | yes |
| Mikicin et al., 2015 |  | 35 | C3 and C4 electrodes | 18-25 yrs |  |  | yes |
| Jurewicz et al., 2018 | AAT – anticipatory attention test | 32 | F3, F4, P3, and P4 | 22.34 ± 1.18 | move points to center of target | up versus downregulation | no |
| Studer et al., 2014 | attention network test | 55 | Cz | 19-31 | changing bars, ball position | two feedback groups plus control cognitive training (not sham) | no |
| Mirifar et al., 2019 | simple and choice RT | 38 | Cz | 16.8 (SD 2.47) | light bulb, with score and beeps | two feedback groups plus sham control group | no |
| Tseng et al., 2021 | episodic and semantic memory encoding and recall | 27 | Fz | 21.6 years, SD = 4.15 | pitch of a tone | no-NFT passive control | yes |
| Ghaziri et al., 2013 | “attentional performance” via the Integrated Visual Auditory (IVA) continuous performance test | 30 (12 exp, 12 sham, 6 control) | F4, P4 | 22 (2.4) | 2 columns, green when above thres, red when below | sham with feedback from exp group, and control with no training | yes |
| Cheng et al., 2015 | putting performance (distance from hole in cm) | 16 (8 exp, 8 sham) | Cz | exp: 22.3 (2.07), sham: 20.6 (1.59) | graphical feedback representations including the low-frequency audio-feedback tone by acoustic bass | sham group | yes |

Many of the included manuscripts did not report all statistical information needed to estimate the effect size. Where this information was available, the algorithm proposed by Rosnow et al. (1996) was utilized to determine the effect size measured as standardized mean differences, i.e., as Cohen’s D. This method can be used on t-tests, means, and standard deviations, as well as F-tests with 1 degree of freedom. As previously noted in the review by Rogala et al. (2016), statistical models for establishing outcomes of NFT vary. Some include (1) mixed model ANOVA, comparing active and control groups at pre- and post-intervention, with an interaction between group and pre-post a hallmark of effects of NFT on behavior; (2) between-group t-tests at post; (3) within-participants t-tests within active and control group, and interpreting significant effects in active versus control as evidence of NFT having an effect. Many journal guidelines recommend not to use strategy (3) for defining a group by treatment interaction (Cumming, 2013), but for the present review, these were included for papers using strategy (3), while also calculating effects sizes based on the active-group’s pre to post comparison.

As mentioned above, where multiple dependent variables for a given task were reported, or where multiple tasks existed, the most widely used index of that task (e.g., response time in a response time task) was selected for reporting in Tables 1 and 2, but all tasks and indices entered the descriptive statistical analysis and effect size calculations. Effect sizes were not examined in a meta-analysis because the studies showed a large degree of heterogeneity in terms of paradigm, analyses, and concept/behavioral targets. Effect size distributions and other descriptive statistics are presented instead to characterize central tendencies within subsets of the literature that allow such descriptive comparisons.

## Results

### Search Results

An initial search for neurofeedback articles retrieved 328 records from PubMed and 23 records from the two other databases. Of these, 264 (108 fMRI and 156 EEG) articles were excluded due to (1) No neurofeedback task (e.g., BCI study), (2) No behavioral outcome, (3) No control group or control condition, (4) A review study, and/or (5) Clinical or developmental populations. Further, 23 (6 fMRI and 17 EEG) articles were excluded due to small sample size (N < 4) and/or a lack of reporting any statistical information on the behavioral outcomes. The final set of studies included in the present report in Tables 1 and 2 comprised 16 studies with fMRI and 48 with EEG. Figure 2 shows their distribution across publication years illustrating a recent trend toward seeing more studies measuring behavioral outcomes. This may be a consequence of an increasing base rate of published NFT studies as noted in recent review papers (Rogala et al., 2016).

**Figure 2.**
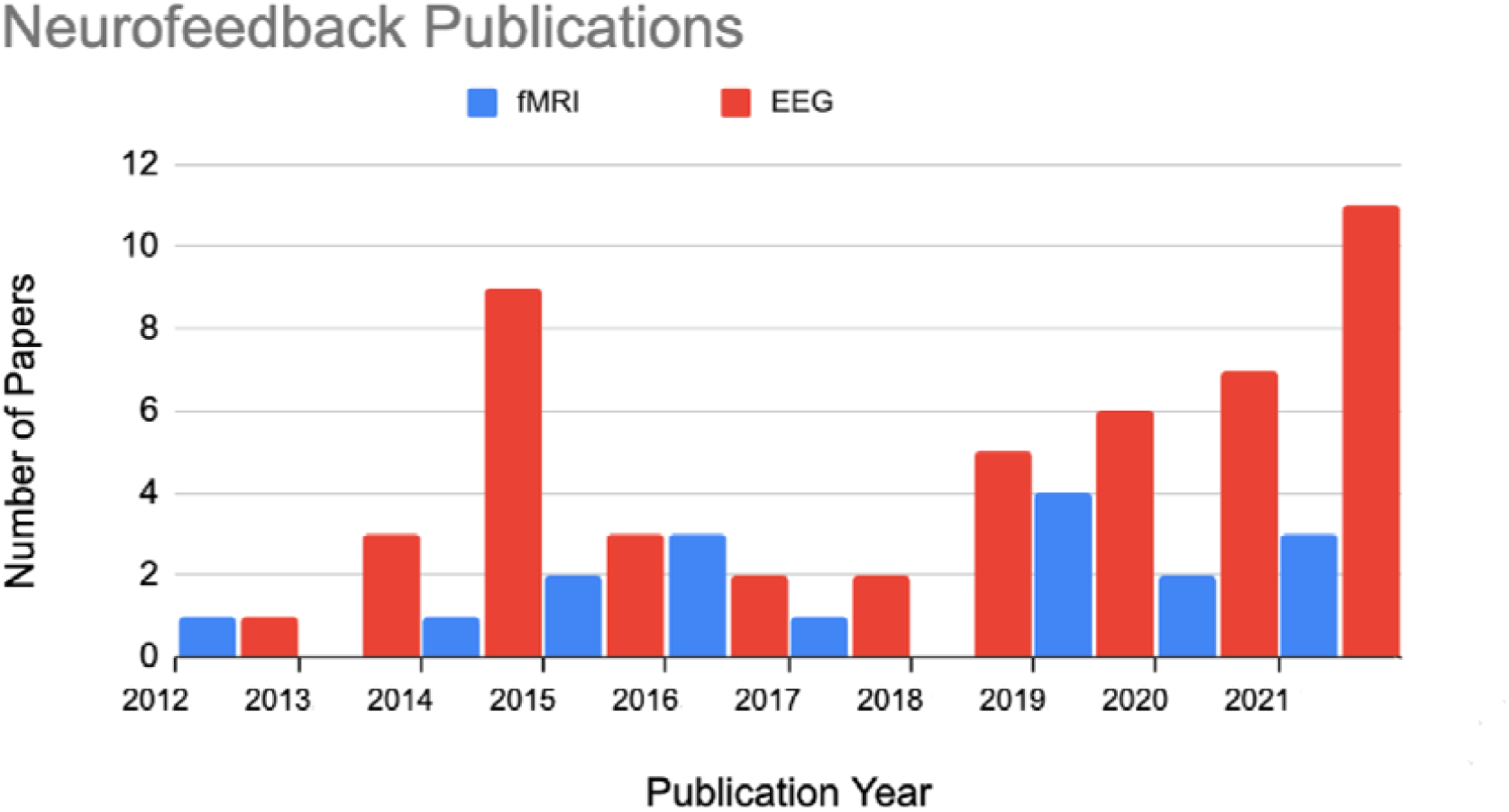
Count of studies, by publication year, for fMRI and EEG neurofeedback studies included in this PRISMA review.

### Sample Size

Studies varied widely in terms of targeted neural processes, behaviors, study design, and analytical models. This includes substantial variability in terms of sample size. Studies with very small samples were excluded from this review, but many of the included studies would likely still be seen as based on an insufficient number of observations by many statisticians (Brysbaert, 2019). Figure 3 shows the frequency of different sample sizes being used across all included studies. The included studies are summarized in Tables 1 and 2, separately for fMRI (n = 16 studies) and EEG (n = 49 studies).

**Figure 3.**
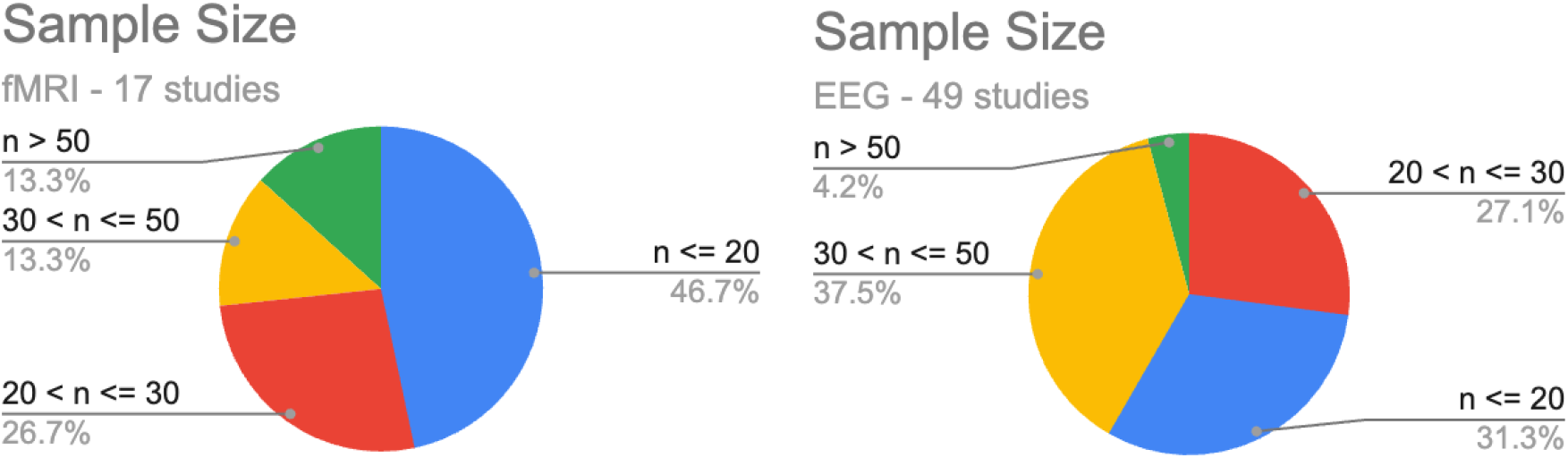
Proportions of studies with different sample sizes for both fMRI and EEG studies included in the present analyses. Note that most studies used samples of 30 participants or less.

**Figure 4.**
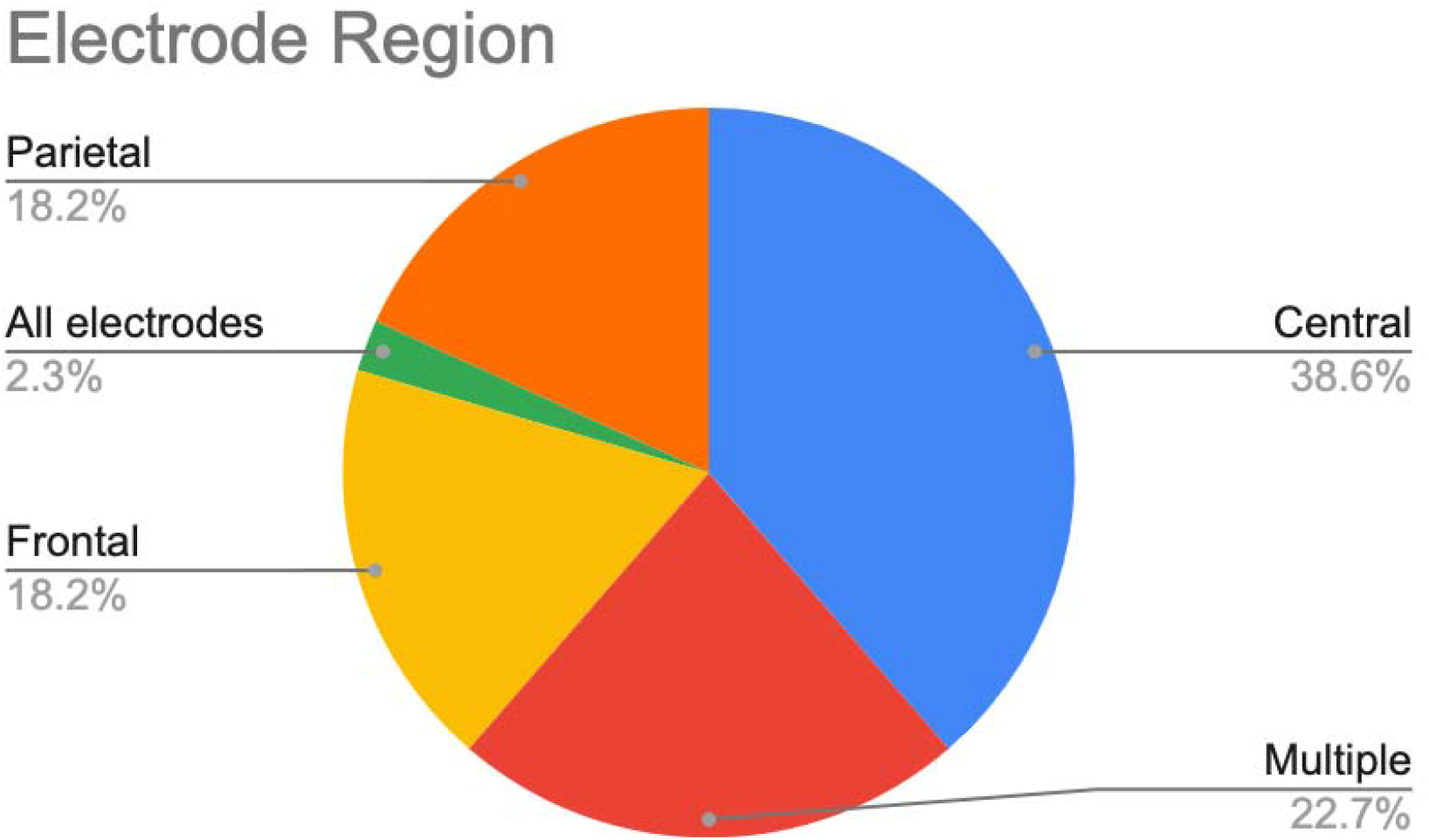
Proportions of studies targeting different aspects of brain activity in EEG-based NFT.

Across all studies selected, sample size (see Figure 3) was below recommended standards for human fMRI or EEG studies (Desmond & Glover, 2002; Keil et al., 2014), given previously reported effect sizes (Gruzelier, 2014a; Rogala et al., 2016). While EEG studies indicated approximately 42% of studies with a total sample size of n > 30, 29% of fMRI studies reported an n > 30, often split between more than two groups of participants. In a similar vein, 46.7% of fMRI studies and 31.3% of EEG studies had sample sizes below 20%.

### Study Design

Across all 65 articles selected, about 50 studies chose an active control group and condition design (sham stimulation, yoked to another participant, etc.) over a passive control (e.g., waiting list, behavioral training only). The sensory modality of the feedback representing the targeted brain activity was mostly visual (53.6%). A minority of studies employed a combination of modalities (5.4%).

### EEG NFT targets: Electrode location and frequency

Across all 49 EEG studies selected, researchers focused on analyzing activity at central electrodes. Specifically, sensors C3, Cz, and C4 were most frequently used, reflective of significant interest in NFT on motor behaviors seen across many studies included in this review.

In terms of EEG band power frequency, as shown in Figure 5, brain activity in the canonical alpha frequency band (8 – 13 Hz) was most frequently targeted (34.9%). Brain activity in the Gamma frequency band (> 20 Hz) was the least frequently used (2.3%). For NFT on ratios between two frequency bands, theta and beta rhythms tended to be used the most (i.e., theta/beta ratio).

**Figure 5.**
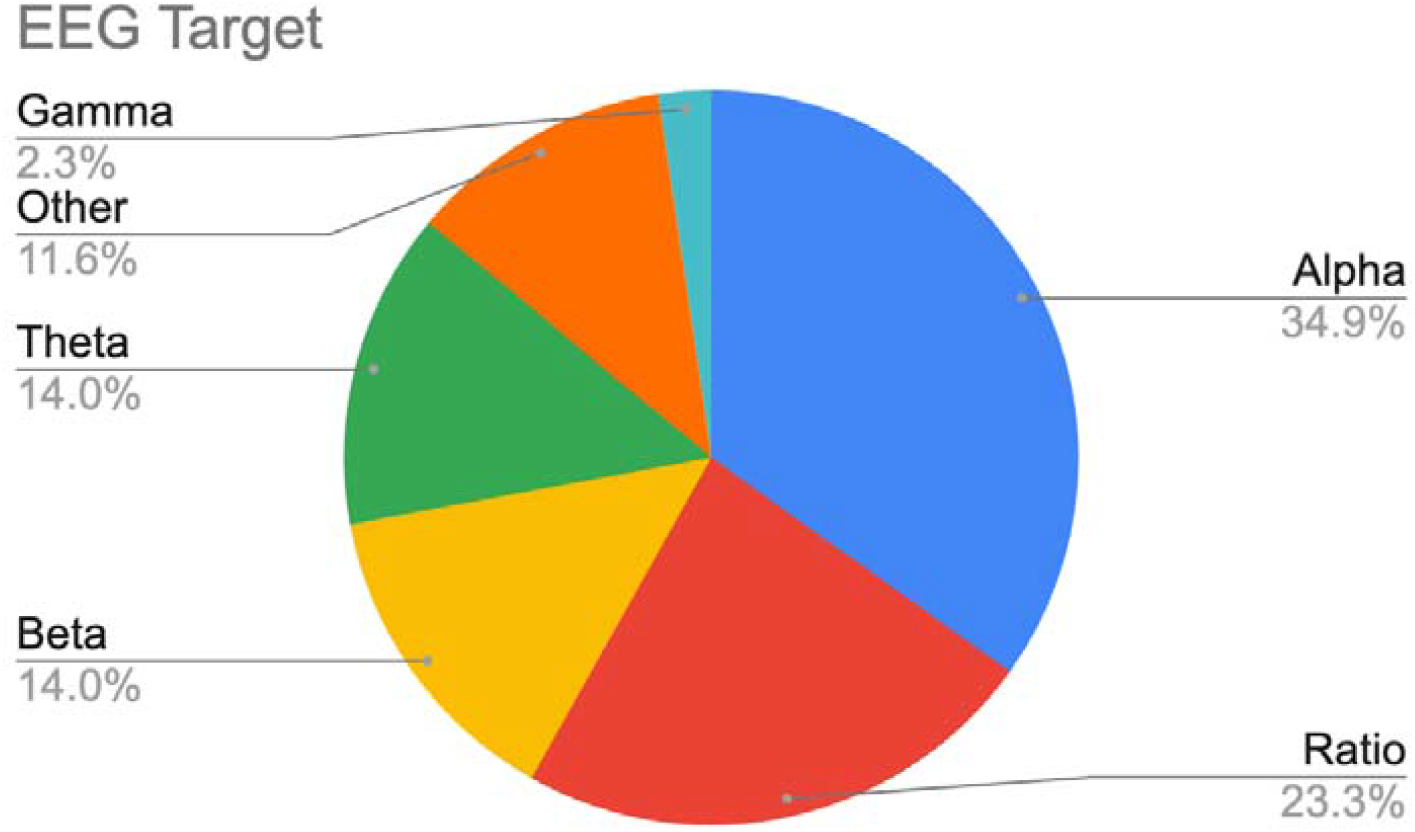
Proportion of NFT studies targeting different canonical frequency bands.

### Effect Size and Frequency of Significant Effects of NFT on behavioral outcomes

When considering the p-values of all statistical tests reported on all behavioral outcomes, across all fMRI and EEG NFT studies, only 44% of these tests reported a significant effect. As shown in Figure 6, the same overall pattern emerges when considering the effects at the study level rather than at the level of individual tests across all studies: Because many of the included studies employed multiple tasks and, therefore, reported multiple p-values, results were counted significant if at least 50% of p-values for each task showed significance, as defined liberally (p<.05). When this quantification was applied, 58.8% studies with fMRI and 47.9% of studies with EEG reported a preponderance of absence of effects.

**Figure 6.**
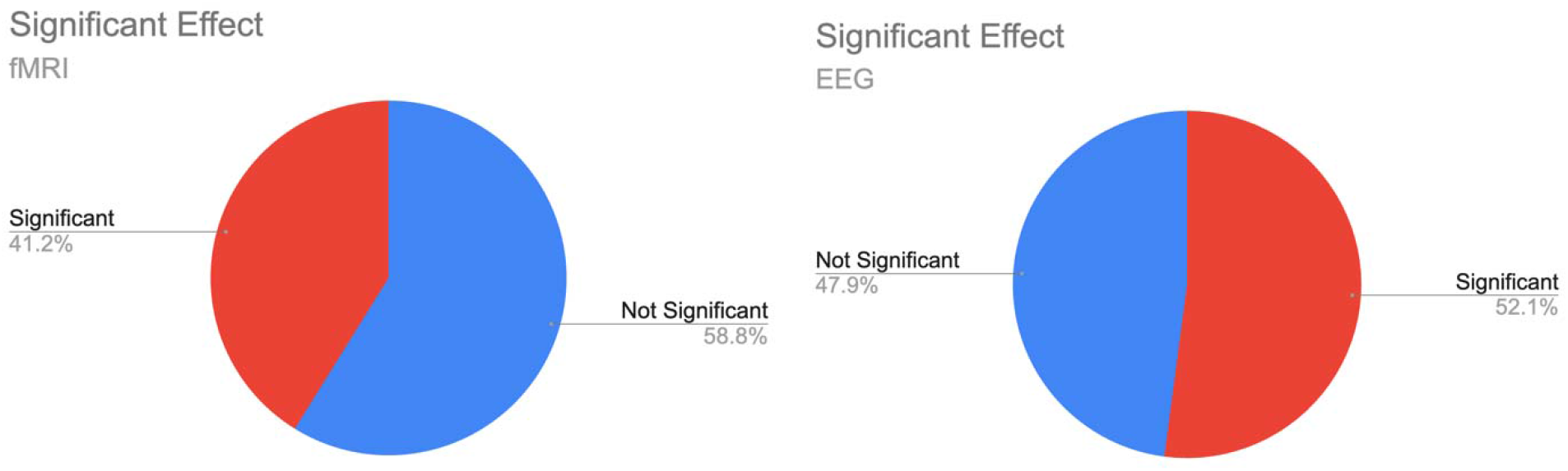
Pie charts of significant and non-significant reported p-values for fMRI and EEG studies included in analyses, quantified at the level of study. In cases where more than one dependent variable was reported without the original authors denoting a particular variable as the central outcome, a result was counted as “significant” in which most outcomes showed statistically significant (p<0.05) results.

Effect sizes (Cohen’s d, Rosnow et al., 1996) were available or could be estimated for 13 (out of 16) fMRI studies and 40 (out of 48) EEG studies. Effect size analyses mirrored the results of the thresholded outcome analysis above. As illustrated in the histograms in Figure 7, although several studies reported medium to large effects, most reported small and medium effects. Median estimated effect size, unweighted by sample size was .54 for fMRI, and .43 for EEG, consistent with medium-sized effects (Rosnow et al., 2000). However, when linear weighting was applied by the sample size, weighted mean effect size values were reduced (.27 for fMRI and .32 for EEG). Effects in the opposite direction as expected were briefly mentioned in several studies but tended to be unaccompanied by statistical indices. Thus, these effects are not included here. In summary, although there was a substantial subset of studies with what appeared to be robust effects of NFT on behavioral outcomes, most studies showed very small or absent effects. To explore predictors of effect size, non-parametric (Spearman’s rho) correlations were calculated between effect size estimates and several potential predictors, including sample size and number of training sessions. Across all studies with effect sizes, there was no relation between training session count and effect size (rho = -.24, p = .10) but there was a strong negative relation between sample size and effect size (rho = -.39, p = .005). Finally, crosstabulation analyses linking significant/non-significant outcomes to targeted frequencies (*χ*^2^[5] = 5.27, p = .39), and to the type of control condition or control group (*χ*^2^[1] = 0.022, p = .88), showed no such association. Rates of significant findings were not more likely in EEG than in fMRI studies (*χ*^2^[1] = 1.02, p = .31).

**Figure 7.**
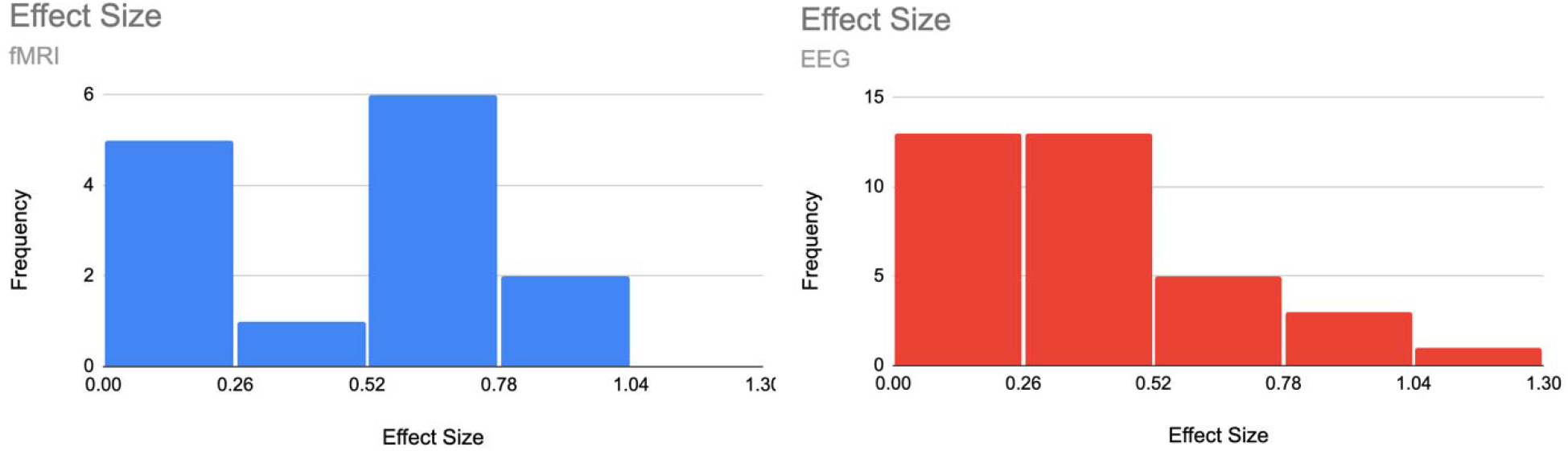
Histograms of the effect sizes for both fMRI (left panel) and EEG studies (right panel) included in analyses. Note that effect sizes were not available for all studies included but only for a subset of studies.

## Discussion

The aim of this systematic review was to examine the effects of a wide range of NFT interventions on outcomes in experimental tasks with dependent variables operationally defined by a task, such as accuracy or response time, in non-clinical individuals. Clearly defined affective outcomes such as evaluative ratings of valence and arousal were also included. Considering studies using both fMRI and EEG, overall weak evidence for NFT’s effect on task performance was found.

Regarding NFT’s effectiveness on outcomes in experimental tasks, results show that less than 50% of studies reported performance improvement with NFT. Pooled and weighted effect sizes of Cohen’s ds (.27 for fMRI and .32 for EEG respectively) also suggest that although several studies reported strong effects, the entirety of studies considered here did not show strong effects of NFT on task performance. As did prior reviews with conclusions similar to ours (Rogala et al., 2016), the present findings differ from the more positive assessment of effect sizes as given in earlier reviews targeting healthy populations, for example in Gruzelier (2014a, b) and in a recent systematic review and meta-analysis focusing on athletes’ decision making and reaction times by Brito et al. (2022). One source of discrepancy between the present and earlier findings may be rooted in the many reviews, discussions, and recommendation articles that have been published in recent years (Arns et al., 2017; Schabus, 2017; Taschereau-Dumouchel et al., 2021), and which may have changed practice in the field. For example, prior recommendations to use active control groups may have impacted the field and thus may have led to more robust estimations of small effects, compared to earlier work with less appropriate experimental control (Rogala et al., 2016). In addition, the disposition of authors and journals towards publishing null or negative findings has changed in the past decade, even though present analyses did not reveal any registered reports and few preregistrations among the studies included in this review. The difference between present findings and those of Brito et al. may be attributed to the different studies included in their meta-analysis, compared to the present review.

Overall, paralleling earlier review articles, present results show that studies varied widely in sample size, data processing, artifact control, and, importantly, research design. Despite earlier recommendations, several studies used passive control conditions discouraged in previous review papers (Rogala et al., 2016), and only a minority used double-blind randomized cross-over designs, often considered the most appropriate research design for studying intervention effects (Schabus, 2017). Furthermore, reporting of statistical indices needed for meta-analytical evaluation is variable in this literature. Future work may wish to report means and variability for each condition and group, along with full inference statistics. Sample sizes were overall smaller than widely recommended in studies addressing cognitive neuroscience questions, with many studies featuring cell sizes of 15 and below. Given the intervention character of NFT studies, basing sample sizes on suitable power analyses, including simulations that include trial counts (Boudewyn et al., 2018; Gibney et al., 2020), may provide an avenue toward more robust, converging findings. Notably, there was a negative correlation between effect size and sample size, suggesting that small-sample studies overestimate effect sizes, a known issue that has been widely discussed in biobehavioral research (Begley, 2013; Halsey et al., 2015; Larson, 2020). There was no association between finding a significant effect of NFT and NFT modality (EEG, fMRI), targeted brain process, or study design. The lack of behavioral outcomes in many studies may also inform other procedures used in NFT research: Many studies use a brain-behavior metric, sometimes based on correlating changes in the brain with changes in behavior. If there is a null effect of NFT on behavior however, any NFT-related differences observed on these measures might be driven by differences in brain activity only. In a similar vein, many of the NFT research papers considered here did not specify whether the goal of the intervention is to prompt behavior change, or brain activity change, or change in various combined brain-behavior metrics. Future work may use systematic manipulation and more refined meta-analytic categorization to examine potential effects of different experimental paradigms and different analytical approaches on NFT efficiency.

Given the strong reliance of EEG-NFT on band power measures derived through spectral analysis, authors working in NFT research may wish to consider recent developments in spectral decomposition of neural time series (Donoghue et al., 2020; He, 2014). These advancements have led the field to consider problematic the process of extracting band power averages from spectra without considering potential changes or group differences in the overall spectral shape (Keil et al., 2022). Many suitable algorithms are available for avoiding confounds of specific activity in a given frequency band and overall spectral shape (Barry & Blasio, 2021; Donoghue et al., 2020; Hughes et al., 2012). NFT researchers may wish to examine the value of these approaches for heightening internal and external validity of EEG-based NFT. In a similar vein, fMRI-based NFT has increasingly relied on multivariate decoding methods (Taschereau-Dumouchel et al., 2021), and the potential of these advanced methods has shown promise for basic science and applied research alike (Cortese et al., 2021; Z. Wang et al., 2021a).

Finally, how long the effect of NFT remains, at both neural and behavioral levels, after terminating an intervention is also a topic for future investigation. NFT studies are often designed and tested with immediate post-test in mind. Potential advantages of including short-term and even long-term follow-up measurements in NFT studies are: a) the ability to examine the temporal evolution and persistence of effects; b) the ability to quantify the reliability and variability of effects over time; and c) the ability to quantify lasting real-world benefits of the intervention.

In summary, the present PRISMA review converges with earlier reports (Rogala et al., 2016), showing that NFT research in normative, healthy samples is heterogenous regarding key methodological issues. Substantial variability was found in terms of sample size, experimental control, data processing, targeted brain processes, and statistical methods, among other criteria. Much of this variability likely contributed to variability in outcomes, with only a minority reporting significant effects of NFT on task performance. The present study does not suggest that there are no effects of NFT on behavioral performance in experimental tasks. Instead, paralleling earlier reviews of this literature (Rogala et al., 2016), the present analysis suggests that future reports in NFT research may wish to emphasize power analyses, pre-registration, and registered reports, as well as fully and carefully reporting statistical metrics needed for assessing the robustness of findings. Bayesian statistical approaches may also be helpful when aiming to quantify the meaningfulness of null findings. Lastly, NFT studies will be more persuasive once researchers more widely engage in direct replication research and include long-term follow-up in their study designs.

